# Mutual transcriptional repression between *Gli3* and *Hox13* genes determines the anterior-posterior asymmetry of the autopod

**DOI:** 10.1101/419606

**Authors:** M^a^ Félix Bastida, Rocío Pérez-Gómez, Anna Trofka, Rushikesh Sheth, H. Scott Stadler, Susan Mackem, Marian A. Ros

## Abstract

In the present study we have investigated the molecular causes of the absence of digit 1 in the *Hoxa13* mutant and why the absence of Hoxa13 protein, whose expression spans the entire autopod, specifically impacts the anterior-most digit. We show that in the absence of *Hoxa13*, the expression of *Hoxd13* does not extend into the anterior mesoderm consequently leaving the presumptive territory of digit1 devoid of distal *Hox* expression and providing an explanation for the agenesis of digit 1. We provide compelling evidence that the lack of *Hoxd13* transcription in the anterior mesoderm is due to increased Gli3R activity, in turn resulting from the loss of transcriptional repression exerted by Hoxa13 on *Gli3*. Our results are compatible with a mutual transcriptional repression between *Gli3* and *Hox13* genes that determines the anterior-posterior asymmetry of the autopod.

## INTRODUCTION

The developing vertebrate limb has long proved as an excellent system for studying the mechanisms involved in pattern regulation and morphogenesis (Zeller et al., 2009) (Zuniga, 2015). Many of the genes important for limb patterning have been identified but little is known about the mechanistic implementation of gene expression patterns into specific morphological traits.

Among the genes essential for the outgrowth and patterning of the tetrapod limb are the *Hox* genes (Zakany and Duboule, 2007). Members of the *HoxA* and *HoxD* clusters display complex and dynamic patterns of expression during limb development that contribute to the organization of limb morphology (Spitz et al., 2001; Tarchini and Duboule, 2006; Zakany and Duboule, 2007).

A large body of work has established that the expression of *Hoxd* genes takes place in two successive phases. The first phase occurs in the emerging limb bud, principally involves expression of *Hoxd8* to *Hoxd11*, and correlates with the specification of the upper-arm (stylopod) and forearm (zeugopod) morphology (Tarchini and Duboule, 2006; Woltering and Duboule, 2010). The second phase of *Hoxd* gene expression occurs in the hand plate, mainly involves *Hoxd10* to *Hoxd13*, and is associated with the morphology of the hand (autopod) (Kmita et al., 2002; Spitz et al., 2001; Woltering and Duboule, 2010). The domains of expression corresponding to each of the two phases of expression are clearly separated by a transversal band of tissue, at the prospective wrist/ankle level, devoid of *Hoxd* transcripts (Woltering and Duboule, 2010).

Recent investigations have revealed that these precise patterns of expression rely on complex transcriptional regulation that involves multiple long-range enhancers located within the flanking topologically associating domain (TAD), regions of the chromatin with a discrete three-dimensional architecture in which internal interactions are favored (Andrey et al., 2013; Dixon et al., 2012; Lonfat et al., 2014). First phase transcription relies on enhancers located telomeric to the cluster, in the so-called T-DOM region, while the second phase transcription relies on enhancers located centromeric to the cluster, in the so-called C-DOM region (Andrey et al., 2013; Montavon and Duboule, 2013; Montavon et al., 2011). Hoxa13 has recently emerged as a major regulator of the switch between these two types of transcriptional regulation terminating the T-DOM dependent proximal regulation and re-enforcing the C-DOM distal regulation (Beccari et al., 2016; Sheth et al., 2016).

The *HoxA* cluster is the other Hox cluster involved in instructing limb morphology with 5’ *Hoxa* genes sequentially activated in the distal limb bud (Boulet and Capecchi, 2004; Davis et al., 1995; Fromental-Ramain et al., 1996; Kmita et al., 2005). Similarly to *Hoxd* genes, the transcription of *Hoxa* genes depends on remote enhancers scattered over the genomic landscape upstream of the cluster (Berlivet et al., 2013). Interestingly, the eventual activation of *Hoxa13* in autopod progenitor cells abrogates *Hoxa11* expression, generating mutually exclusive *Hoxa11-Hoxa13* domains of expression that define the two distal segments of the limb, the zeugopod and the autopod respectively (Tabin and Wolpert, 2007). The activation of *Hoxa13,* together with *Hoxd13*, in the autopod progenitors controls expression of *Hoxa11* sense and antisense transcription in a negative and positive way, respectively (Kherdjemil et al., 2016; Sheth et al., 2014; Woltering et al., 2014). Thus, Hoxa13 is a major regulator of both *Hoxd* and *Hoxa* gene expression.

Another major regulator of *Hoxd* gene transcription is Gli3, the principal transducer of Sonic hedgehog (Shh) signaling during limb development (Hui and Joyner, 1993; Litingtung *et al.*, 2002; Lopez-Rios, 2016; Schimmang *et al.*, 1992; te Welscher *et al.*, 2002). In the absence of Shh, Gli3 is processed to a short form that acts as a strong transcriptional repressor (GLI3R) (Wang et al., 2000). Initially Hox proteins contribute to activate *Shh* transcription (Capellini et al., 2006; Kmita et al., 2005) but then Shh function is essential for the second phase of *Hoxd* expression by relieving the Gli3R repression (Lewandowski et al., 2015; Vokes et al., 2008). In the absence of *Gli3*, as in the *extratoes* (*Xt*) spontaneous mutation in mice, the autopod is characterized by a prominent uniform anterior-posterior (AP) expansion of *Hoxd* expression without a noticeable change in *Shh* expression (Zuniga and Zeller, 1999).

In addition to their temporospatial transcriptional control, the function of Hox products can also be modulated by interaction with co-factors. Some Hox products have been shown to interact with other DNA-binding co-factors including Smad5, Gli3 and Hand2 (Chen et al., 2004; Galli et al., 2010; Williams et al., 2005). For example, through physical binding, Hoxd12 can sequester Gli3 repressor (Gli3R) and even at high levels convert it from a transcriptional repressor to a transcriptional activator contributing in this way to the regulation of digit patterning (Chen et al., 2004). Likewise, proteins of the Hox13 paralogous group can modulate Bmp and TGFβ/Activin signaling activity through protein-protein interaction with Smads (Williams et al., 2005) and associate with Hand2 to activate *Shh* transcription (Galli et al., 2010).

*Hoxa13* is the only member of the 39 mammalian *Hox* genes whose deletion is embryonic lethal as it is required for proper placental function (Scotti and Kmita, 2012; Shaut et al., 2008). Here we have focused on the study of the *Hoxa13*^*-/-*^ null limb phenotype which is restricted to the autopod and characterized by the absence of the anterior-most digit 1, syndactyly and brachydactyly (Burke et al., 1995; Fromental-Ramain et al., 1996; Goodman et al., 2000; Innis et al., 2002; Mitsubuchi and Endo, 2006; Perez et al., 2010; Stadler et al., 2001). Because of the importance of digit 1, the thumb in the human hand, we asked why the absence of Hoxa13 protein, whose expression spans the entire autopod, specifically impacts the anterior-most digit. In contrast, although Hox paralogs often display considerable functional overlap, selective *Hoxd13* loss does not impair digit 1 formation (Dolle et al., 1993). We report that in the absence of *Hoxa13*, the expression of *Hoxd13* does not extend into the anterior mesoderm consequently leaving the presumptive territory of digit1 devoid of distal *Hox* expression, a circumstance that is considered sufficient to prevent digit condensation (Fromental-Ramain et al., 1996; Sheth et al., 2012). We also show that the lack of *Hoxd13* transcription in the anterior mesoderm of *Hoxa13* mutants correlates with increased Gli3R activity resulting from the loss of the transcriptional repression of *Gli3* exerted by Hoxa13. Our results are compatible with a mutual antagonism between Gli3 and Hox13 genes that determines the anterior-posterior asymmetry of the autopod.

## RESULTS

### A gene dosage effect of *Hoxa13* in digit 1 morphology

*Hoxa13*^*-/-*^ mutants die at mid gestation, usually between E12.5 and E14.5, due to placental and vascular defects (Fromental-Ramain et al., 1996; Scotti and Kmita, 2012; Shaut et al., 2008; Stadler et al., 2001). Although it has been reported that a small percentage of *Hoxa13* homozygous mutants survive to adulthood in the C57BL/6J genetic background (Perez et al., 2010), no homozygous pup was born in our colony despite being maintained in this genetic background. Indeed, the oldest homozygous embryos that we recovered were at embryonic day 16.5 (E16.5). At this stage, the typical *Hoxa13* null limb phenotype, consisting predominantly of absence of digit 1 and syndactyly, was prominent (Fig. 1A; (Fromental-Ramain et al., 1996; Stadler et al., 2001)).

**Figure 1.**
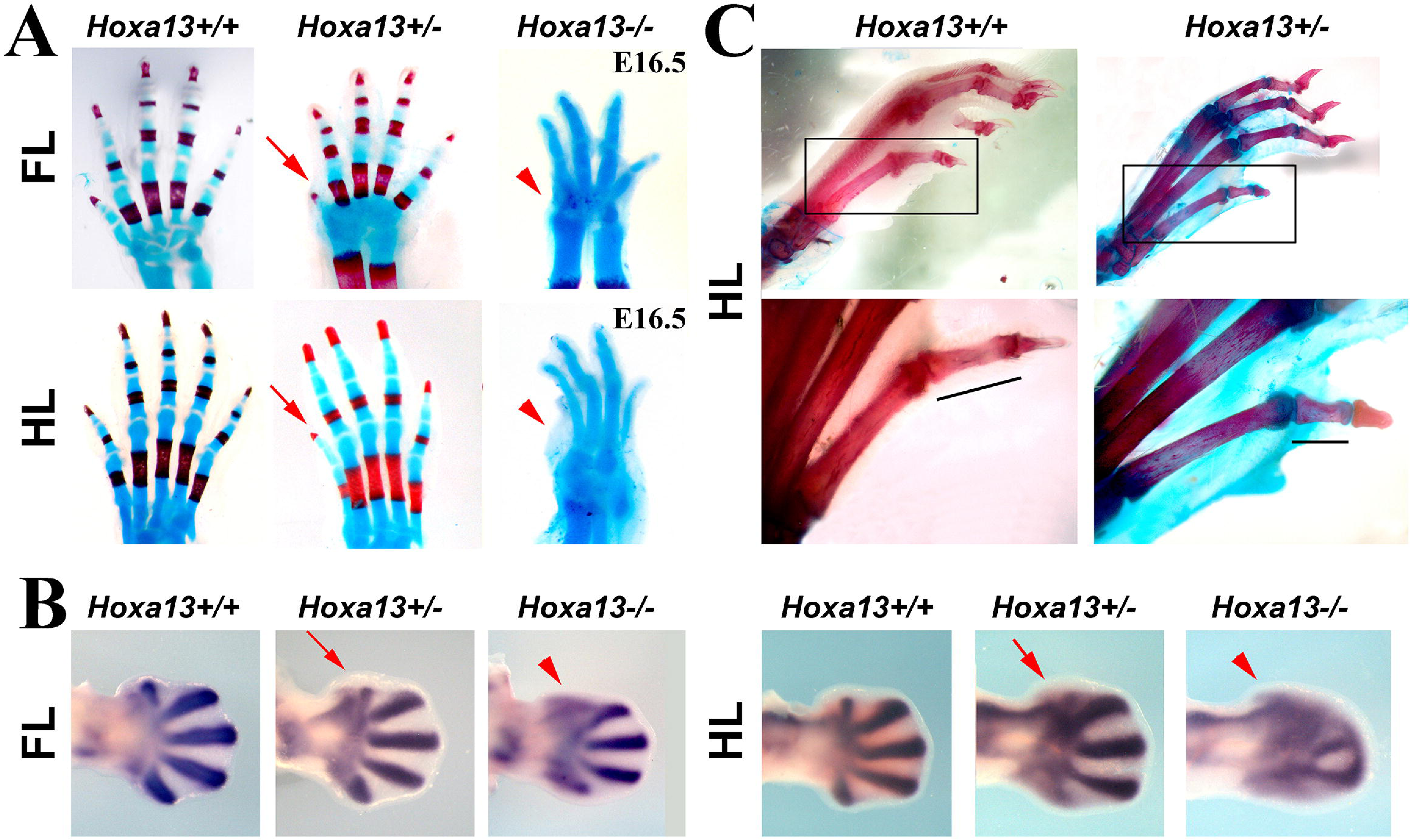
*Hoxa13* mutant limb phenotype. **A)** Alcian blue-alizarin red staining of wild type, *Hoxa13* heterozygous and *Hoxa13* homozygous mutants fore and hind limbs at the stages indicated at the panel top. **B)** Expression of *Sox9* in the E12.5 autopod of wild type, *Hoxa13* heterozygous and *Hoxa13* homozygous fore and hind limbs. **C)** Skeletal staining of 3-month-old wild type and *Hoxa13* heterozygous littermates indicating the reduction in length of the first phalanx of digit 1 (framed area magnified below). Arrowheads and arrows point to the digit 1 phenotype of *Hoxa13* null and heterozygous mutants respectively.

A careful inspection to determine the onset of the limb phenotype showed that the external aspect and shape of the mutant limb was indistinguishable from normal up to E11-E11.5. This was expected since activation of *Hoxa13* normally occurs at E10.5. By E12.5, a flattening of the anterior border of the mutant autopod became progressively conspicuous reflecting the failure of digit 1 to form (Fig. 1B (Fromental-Ramain et al., 1996)). Accordingly, the expression of *Sox9*, the best marker of chondroprogenitors and differentiated chondrocytes, remained diffuse in the prospective digit 1 area without evidence of a digital condensation (arrowhead in Fig. 1B). In heterozygous embryos, the condensation corresponding to digit 1 was less well defined than in wild type littermates (arrow in Fig. 1B) supporting a gene dosage effect for *Hoxa13*. We also noticed that adult heterozygotes showed, in addition to the already reported partial syndactyly and claw alterations (Fromental-Ramain et al., 1996), a mild but consistent hypoplasia of the first digit, particularly conspicuous in the proximal phalanx of the hindlimb (33% reduction compared with wild type n=6; Fig. 1C). This trait was already observed in newborn heterozygotes (arrow in Fig. 1A).

### Altered *Hox* code expression in the anterior limb bud mesoderm of *Hoxa13*^*-/-*^ mutants

The hallmark of digit 1 is the expression of *Hoxd13* but not the other *5’Hoxd* genes *Hoxd10, Hoxd11* and *Hoxd12*. This unique combination of Hoxd products is achieved during the second phase of *Hoxd* transcription when the expression of *Hoxd13* spreads into the anterior mesoderm of the handplate surpassing the anterior limit of *Hoxd11* and *Hoxd12* domains (Kmita et al., 2002; Montavon et al., 2008). Interestingly, it has recently been shown that *Hoxa13* plays a pivotal role in the transition from phase one to phase two of *Hoxd* gene expression (Beccari et al., 2016; Sheth et al., 2016; Sheth et al., 2014). In the absence of *Hoxa13*, the first phase of expression of *Hoxd11* is abnormally prolonged and, consequently, the gap between the two phases becomes distally displaced (Fig. 2A). Since the prolongation of the first phase is more marked anteriorly, the prospective digit 1 cells now reside within the first phase domain of *Hoxd11* expression (arrow in Fig. 2A and (Sheth et al., 2016; Sheth et al., 2014)). This also occurs with the first phase of the more 3’*Hoxd* genes *Hoxd4, Hoxd9* and *Hoxd10*, that extends over the presumptive digit 1 cells and also with *Hoxa11,* a gene typical of the zeugopod (arrows in Supplementary Fig.1;(Sheth et al., 2014)). In contrast, no gross perturbation of the normal *Hoxd12* expression, which extends to the anterior digit 2 border during the second phase of *Hox* gene expression, was detected in *Hoxa13* null autopods (Fig. 2B). Interestingly, *Hoxd13* transcription failed to extend into the most anterior mesoderm and remained in a domain similar to that of *Hoxd12* (arrowhead in Fig. 2C). Thus, digit 1 progenitors in *Hoxa13* mutants express an altered Hox code that corresponds to the zeugopod, rather than to the autopod, as it includes *Hoxa11* and the first phase of *Hoxd* genes but lacks the characteristic expression of *Hoxd13.* The fact that the prospective digit 1 cells in *Hoxa13* null embryos are devoid of both Hoxa13 and Hoxd13 may account for the loss of digit 1 in the Hoxa13 null limb, as mice lacking both Hoxa13 and Hoxd13 exhibit a profound loss of chondrogenic condensations throughout the autopod (Fromental-Ramain et al., 1996; Sheth et al., 2012).

**Figure 2.**
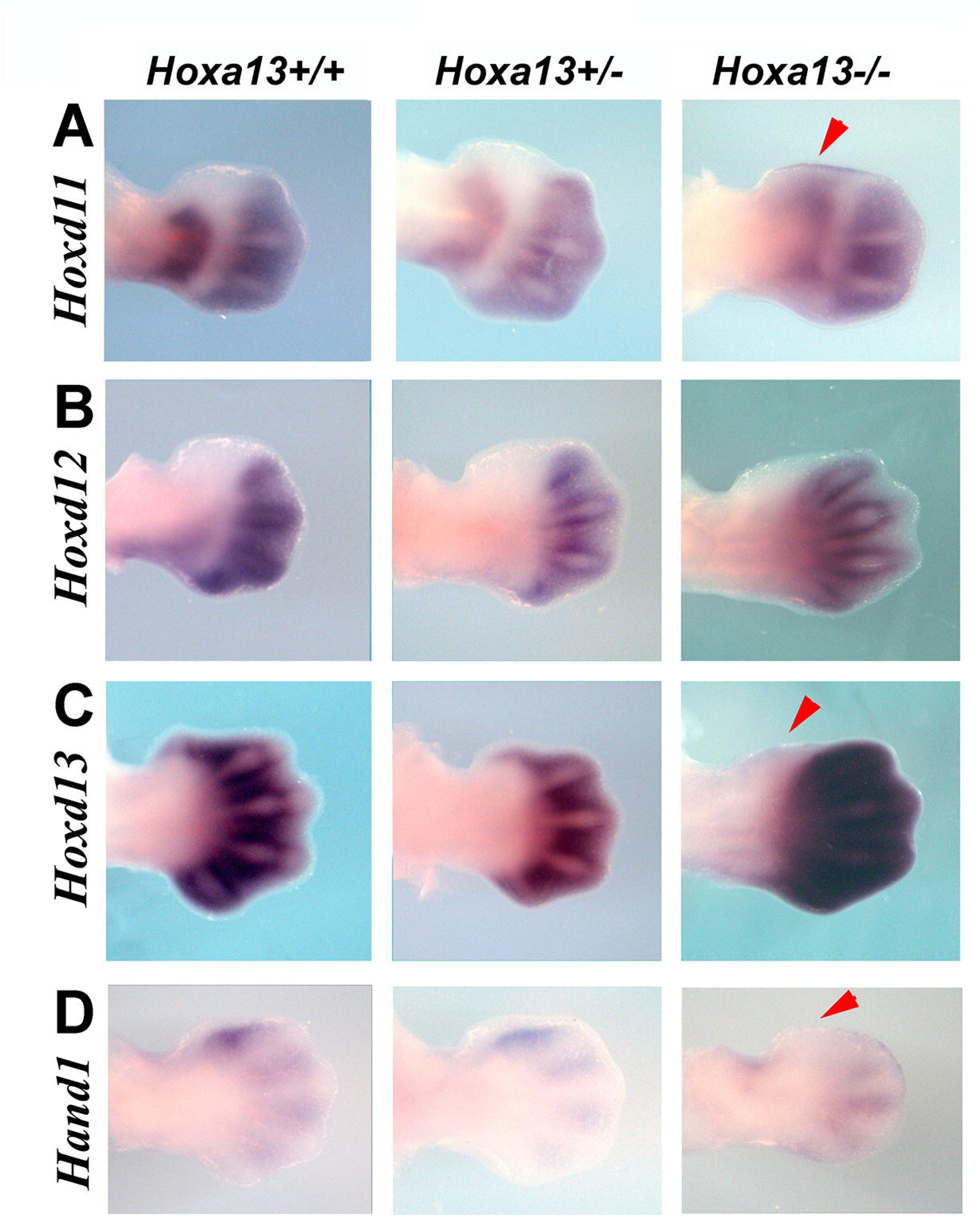
*5’Hoxd* gene expression in *Hoxa13* mutant limb buds. Forelimb autopods of wild type, *Hoxa13* heterozygous and *Hoxa13* homozygous mutants hybridized with *Hoxd11* (**A**), *Hoxd12* (**B**), *Hoxd13* (**C**), and *Hand1* (**D**) at E 12.5. The arrowheads point to altered expression patterns in homozygous mutants.

To further explore the expression of genes characteristic of digit 1 in *Hoxa13* mutants, we also analyzed the expression of *Hand1* (Fig. 2D; (Fernandez-Teran et al., 2003)). In contrast to wild type littermates, *Hand1* expression was not observed in digit 1 in *Hoxa13* mutants confirming the absence of digit 1 specification (arrowhead in Fig. 2D). No differences in the pattern of expression of other markers of the anterior mesoderm, *Tbx2, Tbx3, Alx4*, the expression of which primarily occurs at proximal level, were observed in *Hoxa13* null limb buds compared with wild type littermates (Supplementary Fig. 2). This indicates that the gene expression perturbations are specific of the anterior autopod mesoderm (Knosp et al., 2007).

### No major disturbance of signaling centers in the absence of *Hoxa13*

Since the formation of the digits also depends on the activity of the apical ectodermal ridge (AER) and of the zone of polarizing activity (ZPA), and given the interaction of *Hox* genes with these major signaling centers of the limb bud (Galli et al., 2010; Kmita et al., 2005; Sheth et al., 2013), we decided to investigate the state of the AER and ZPA in *Hoxa13* mutants. *In situ* hybridization at E11.5 showed a downregulation of *Fgf8* in the most anterior AER of homozygous limb buds (arrow in Fig. 3). To assess a possible impact on FGF signal reception, we assayed for the expression of *Dusp6* (formerly *Mkp3*) and *Sprouty4* (*Spry4*), genes considered sensitive readouts of FGF signaling (Minowada et al., 1999; Smith et al., 2006). Reduced *Spry4*, but not *Dusp6,* indicated a minor downregulation of FGF signaling in the territory of digit 1 suggesting that *Spry4* may be a more sensitive readout of FGF signaling than *Dusp6*. *In situ* hybridization for *Shh* and its major targets *Ptc1, Gli1* (Ahn and Joyner, 2004; Harfe et al., 2004) and *Hand2* (te Welscher et al., 2002a), failed to reveal any difference between *Hoxa13* mutants and littermates control limb buds (Fig. 3).

**Figure 3.**
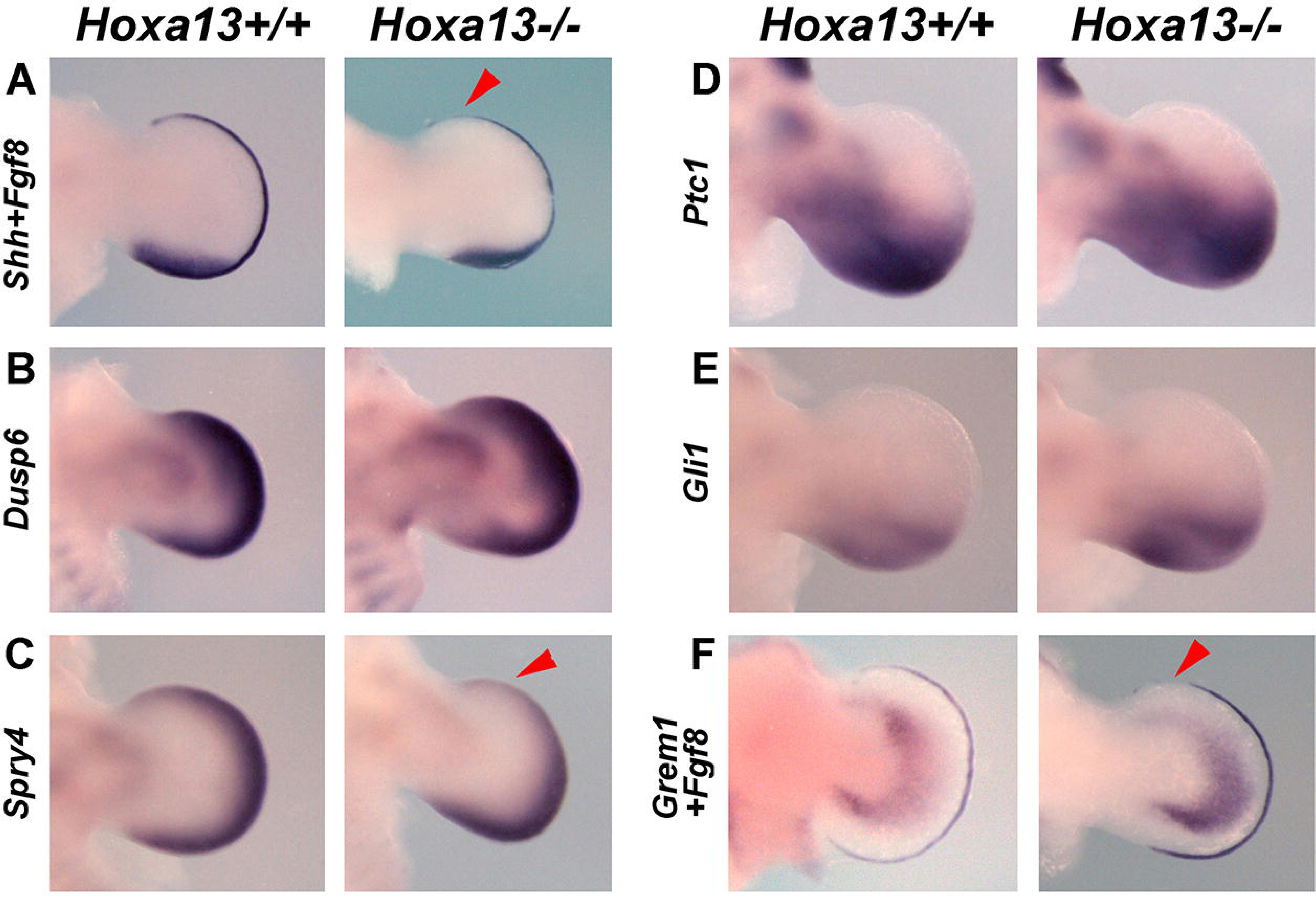
Expression of gene patterning genes in *Hoxa13* mutant limb buds. E11.5 forelimb autopods of wild type and *Hoxa13* homozygous mutants hybridized with *Shh* and *Fgf8* (**A**), *Dusp6* (**B**), *Spry4* (**C**), *Ptc1* (**D**), *Gli1* (**E**), and *Grem1 and Fgf8* (**F**). Note lack of *Grem1* propagation to the anterior mesoderm and reduced anterior *Fgf8* and *Spry4* expression.

Growth during early limb development is supported by a positive regulatory feedback loop established between Shh and AER-FGFs (Bastida et al., 2009; Benazet et al., 2009; Laufer et al., 1994; Niswander et al., 1994; Scherz et al., 2004; Verheyden and Sun, 2008). A crucial component of this regulatory feedback loop is *Gremlin1* (*Grem1*), a Bmp antagonist and Shh target gene, responsible for AER maintenance (Khokha et al., 2003; Michos et al., 2004). Interestingly, the expression of *Grem1* in the *Hoxa13* mutant autopod did not propagate into the anterior mesoderm, as occurs in control limb buds, but remained more posteriorly restricted (Fig. 3). It is likely that the downregulation of *Fgf8* in the anterior AER is secondary to the lack of *Grem1* in the anterior mesoderm. Since Shh expression and signaling are normal, the anterior downregulation of *Grem1* must depend on the alteration of other transcriptional regulators, a plausible candidate being Gli3 (Vokes et al., 2008). Since *Grem1* is primarily regulated by release from repression by Gli3R function, we next wanted to examine the state of Gli3 in *Hoxa13* mutants.

### Increased Gli3R activity in *Hoxa13*^*-/-*^ anterior mesoderm

Our results show that Hoxa13 is required, directly or indirectly, for the normal anterior spread of *Hoxd13* in digit 1 territory. However, we have previously shown that in the double *Hoxa13;Gli3* mutant *Hoxd13* is uniformly expressed at high level all along the anterior-posterior extension of the handplate ((Sheth et al., 2012); Supplementary Fig. 3). Indeed, in the absence of *Gli3* the second phase of expression of all 5’*Hoxd* genes occurs symmetrically all along the anterior-posterior axis of the handplate providing a similar Hox code and presumably a similar amount of Hox products to all digits (Supplementary Fig. 3; (Litingtung et al., 2002; Montavon et al., 2008; te Welscher et al., 2002b)). This suggests that in the absence of *Gli3*, Hoxa13 is no longer necessary for *Hoxd13* transcription in the anterior mesoderm and raises the possibility that Hoxa13 function is required to counteract or modulate Gli3R activity. Therefore, we decided to investigate the expression and activity of Gli3R in *Hoxa13* mutants.

Because Gli3 activity depends on posttranscriptional processing, as a first step to evaluate Gli3R activity we explored the expression of its main target genes. RNA in situ hybridization showed distal expansion of the domains of expression of *Pax9* and *EphA3* two target genes activated by GLI3R (McGlinn et al., 2005), at E11.5 and E12.5 (red arrowheads in Fig. 4A-B). *Bmp4*, a gene whose expression positively correlates with the level of Gli3R (Bastida et al., 2004), was more robustly detected and in an extended domain in the anterior mesoderm (red arrowhead in Fig. 4C). In addition, the GLI3R repressed target gene *Jag1* (McGlinn et al., 2005) was absent from the anterior mesoderm of *Hoxa13* mutants at E11.5 and E12.5 (green arrowheads in Fig. 4D). Overall, the modifications in these expression patterns are consistent with higher Gli3R activity than normal in the anterior mesoderm of the *Hoxa13* mutant. We next wanted to investigate whether this excess occurs at transcriptional or posttranscriptional level.

**Figure 4.**
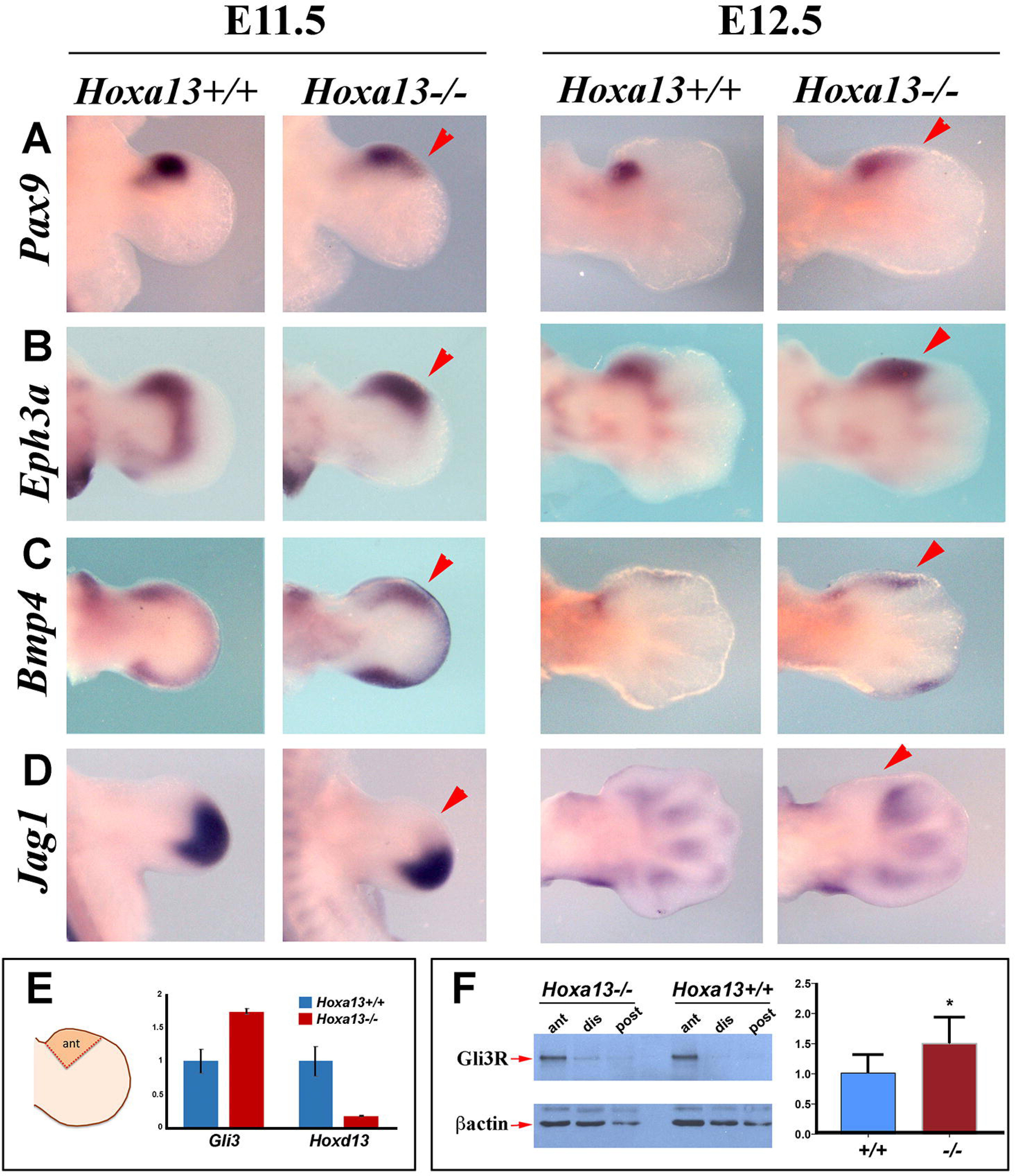
Expression of Gli3R target genes and quantification of *Gli3* mRNA and protein levels in *Hoxa13* mutant limb buds. Pattern of expression of *Pax9* (**A**), *EphA3* (**B**), *Bmp4* (**C**) and *Jag1* (**D**) in E11.5 and E12.5 forelimb buds of wild type and *Hoxa13* homozygous mutants. Altered expressions indicated by arrowheads. RT-PCR quantification of *Gli3* mRNA (**E**) and Gli3R protein (**F**) levels in anterior mesoderm as indicated in the scheme.

To quantify the level of *Gli3* mRNA in the anterior mesoderm of mutants versus wild type, we performed RT-PCR at E11.75. For this we dissected the anterior mesoderm corresponding to digit 1, as depicted in Fig. 4E, the region in which we observed altered gene expressions. Our results showed that the expression of *Gli3* in the *Hoxa13* mutant anterior mesoderm was 1.7 fold higher than in wild type. We also quantified the level of expression of *Hoxd13* by RT-PCR that showed a decrease of 90% confirming our *in situ* hybridization results (Fig. 2C).

To explore Gli3 processing in the mutant, we also used dissected anterior mesoderm fragments of E11.5 forelimbs and analyzed the level of Gli3R by Western Blot (Fig. 4F). Our analysis confirmed a slightly higher level of Gli3R in the anterior limb mesoderm of *Hoxa13* mutant limb buds.

### Altered *Gli3* pattern of expression in the absence of *Hoxa13*

Having shown that the mRNA and protein level as well as the activity of Gli3R, are all increased in the anterior mesoderm, we examined the overall pattern of Gli3 expression in the limb bud. Analysis of *Gli3* expression by in situ hybridization showed a dynamic pattern that has not been previously appreciated. Because of the similarity with the pattern of expression of 5’*Hoxd* genes, we describe it as evolving in two phases (Fig. 5A). As previously described, at E10.5 *Gli3* expression occurred in most of the limb mesoderm except for the most posterior part where *Shh* is expressed (Ahn and Joyner, 2004; Benazet et al., 2009) although the level of expression was less intense in the central mesoderm. By E11, *Gli3* expression in the distal mesoderm became progressively confined to the anterior autopod. By E11.5 a second domain of expression started in the autopod roughly overlapping digit 4 primordium and progressively spanned all the digit primordia (Fig. 5A). Therefore, by E12.5, two domains of expression were clearly distinct and separated by a gap of tissue devoid of transcripts. The proximal domain, remnant of the first phase of expression, remained as a transverse band at the zeugopod-autopod boundary. The distal domain, the second phase of expression, overlapped the distal digit plate and became progressively confined to the interdigital tissues at E12.5 as the digit condensations differentiate (Fig. 5A). By E13.5 the majority of *Gli3* distal expression concentrated in the joints (Fig. 5A).

**Figure 5.**
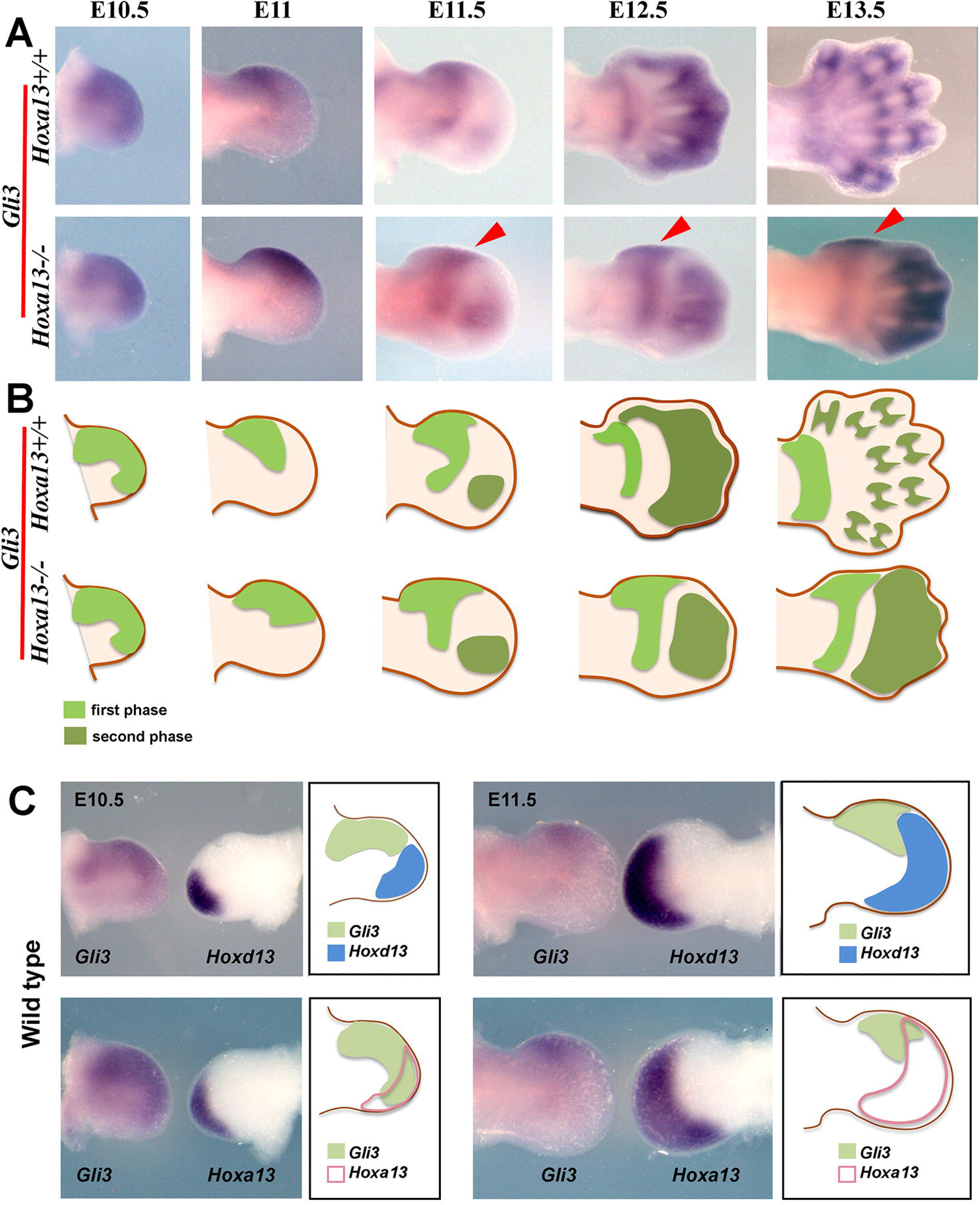
Dynamics of *Gli3* expression pattern in wild type and *Hoxa13* mutant littermates forelimb buds. **A**) Expression of *Gli3* during limb development in control and mutant limb buds. **B**) Schematic representation of *Gli3* expression pattern showing phase one in lighter green and phase two in darker green. **C**) Comparison of *Gli3* and *Hoxd13* and *Gli3* and *Hoxa13* domains of expression in the two limb buds of the same wild type embryo at E10.5 and E11.5. The hybridizations are accompanied by a drawing to better appreciate the relationship between expression domains.

In the absence of *Hoxa13*, the pattern of *Gli3* expression was unaffected up to E10.5 (Fig. 5). However as early as E11, the dynamics of *Gli3* expression were dramatically altered; the downregulation of the first phase of expression was delayed and incomplete, never reaching the digit 1 region (red arrowheads, Fig. 5A). As a consequence, digit 1 territory remained within the first phase of expression of *Gli3* even at later stages, while the second distal domain was restricted to the posterior digits (digits 2-4). A schematic representation of the dynamic expression pattern of *Gli3* and the changes observed in *Hoxa13* mutants is shown in Fig. 5B.

This result uncovered a previously unidentified hierarchical effect of Hoxa13 in *Gli3* regulation. Interestingly, in E12.5 *Hoxa13* mutants the distribution of *Gli3* transcripts was similar to that of *Hoxd10-11* (compare to Fig. 2 and Supplementary Fig. 2) raising the intriguing possibility that Hoxa13 regulates *Gli3* and 5’*Hoxd* genes in a complementary manner. As mentioned in the introduction, Hoxd13 cooperates with Hoxa13 in regulating the switch from the first to the second phase of *Hoxd* gene expression (Beccari et al., 2016; Sheth et al., 2016; Sheth et al., 2014), therefore, we wanted to examine whether *5’Hoxd* genes also played a role in the regulation of *Gli3* expression. In this regard we note that the downregulation of the first phase of *Gli3* transcription correlated with the anterior spread of *Hoxd13* domain. This can be clearly appreciated when the two limbs of the same embryo are hybridized one for *Hoxd13* and the other for *Gli3* and compared (Fig. 5C, and schemes within). The anterior boundary of *Hoxd13* domain coincides with the posterior domain of first phase *Gli3* expression. The comparison between *Gli3* and *Hoxa13* patterns of expression showed overlapping at anterior level that becomes attenuated with time (Fig. 5C).

To start exploring this possibility, we screened the *Gli3* genomic landscape for Hoxa13 and Hoxd13 binding sites using the published data sets of Hoxa13 and Hoxd13 in limb buds, as well as changes in the chromatin state and transcriptome between wild type and *Hoxa13;Hoxd13* double mutants (Sheth et al., 2016). The examination of the *Gli3* locus identified several Hox13 binding regions upstream of the transcriptional start site. Some of these binding regions were also enriched in H3K27ac marks (highlighted in Fig. 6A). Curiously, two peaks (highlighted in yellow in Fig. 6A) overlapped with two previously reported VISTA enhancers (hs1586 and hs1179; (Osterwalder et al., 2018)) with activity in the limb. This result is compatible with a direct regulation of *Gli3* transcription by Hoxa13 and Hoxd13. Our previous in situ and RT-PCR analyses indicated that this regulation was negative and this is confirmed by the increase in *Gli3* transcription (1.7 fold, FDR<= 0.05) that occurs in *Hoxa13;Hoxd13* double mutant autopods, as can be observed in the RNA-seq profiles shown in the bottom tracks in Fig. 6A (Sheth et al., 2016).

**Figure 6.**
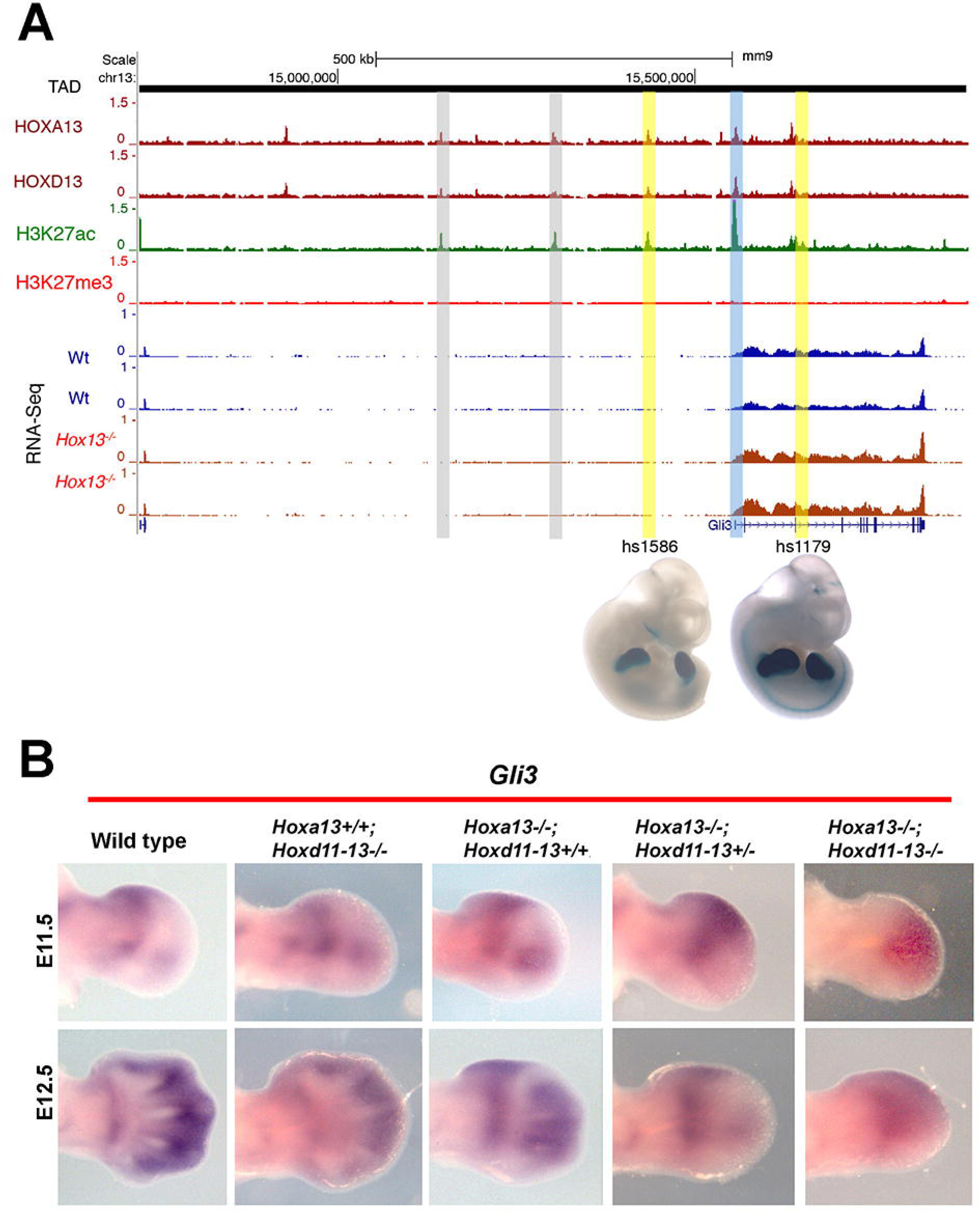
*Hox13* regulation of *Gli3* expression. **A**) UCSC genome browser view of the regulatory landscape upstream *Gli3* (Beccari et al., 2016; Sheth et al., 2016). Black lines indicate the TAD boundaries in ES cells as in Dixon et al. (Dixon et al., 2012). ChIP-seq tracks for HOXA13 and HOXD13 are shown in red on the top. The H3K27ac and H3K27me3 profiles of the digital plate of E11.5 limb buds are also shown in green and red respectively. Two replicates of the transcriptome profiling in wild type (blue) and *Hox13-/-* mutant (red) limb buds at E11.5 are included at the bottom. HOX13 binding sites with potential enhancer activity are highlighted. Two of them, highlighted in yellow, overlap with two previously validated VISTA (Visel et al., 2007) elements (hs1586 and hs1179) and their activity at E11.5 (LacZ reporter, Vista Enhancer Browser) is shown below. **B**) Deregulation of *Gli3* expression in *Hoxa13;Hoxd11-13* compound mutants.

A negative regulation of *Gli3* expression by distal *Hox* genes was confirmed by the changes in expression of *Gli3* in *Hoxd11-13* mutants and *Hoxa13*;*Hoxd11-13* double mutants (Fig. 6B). In agreement with previous results showing that the absence of *Hoxd11-13* had no major impact on *Gli3* mRNA and protein expression (Huang et al., 2016) our analysis additionally confirmed that the pattern of *Gli3* expression was normal in this mutant (Fig. 6B). However, the study of the allelic series showed that the removal of one functional allele of *Hoxd11-13* from the *Hoxa13* homozygous had a stronger impact on *Gli3* expression than the removal of *Hoxa13* alone (Fig. 6B). Finally, in the total absence of distal Hox products (*Hoxa13*;*Hoxd11-13* double homozygous mutants), *Gli3* expression spanned the whole autopod except the posterior border where it is never expressed, again strikingly reproducing the pattern previously reported for *Hoxd10-11* genes in this double mutant (Fig. 6B; (Beccari et al., 2016; Sheth et al., 2014; Woltering et al., 2014)). Interestingly, we have also observed increased *Hoxa13* mRNA expression (unpublished RNAseq, E12.5 autopod, 1.6-fold, FDR=0.01) and protein levels (about 7-fold increase) and protein levels when *Hoxd11-13* gene function was removed (Suppl. Fig. 4). Given the proven redundant function between Hoxa13 and Hoxd13, this increase in Hoxa13 could compensate for the loss of Hoxd11-13 and explain why the removal of *Hoxd11-13* has no effect on *Gli3* transcription (Fig. 6B) or on digit 1 formation.

Together, these results support a dose dependent function of Hoxa13 and 5’Hoxd gene products in regulation of *Gli3* expression mainly controlling the downregulation of its first phase of expression.

### Gli3R repression of *Hand2* operates at later stages

In the emerging limb bud, anterior-posterior patterning is established by the mutual antagonisms between Hand2 and Gli3 (te Welscher et al., 2002a). This antagonism is based on Gli3R directly repressing *Hand2* in the anterior limb bud mesoderm (Vokes et al., 2008) and Hand2 repressing *Gli3* transcription directly and through Tbx3 (Osterwalder et al., 2014). It is known that the requirement of Hand2 to repress *Gli3* transcription is transient, as the removal of *Hand2* after the onset of *Shh* expression, has no consequences (Galli et al., 2010). However, it remains unknown whether Gli3 repression of *Hand2* is also operating at later stages. The analysis of *Hand2* expression in *Hoxa13;Hoxd11-13* compound mutants offers a good opportunity to investigate this question. We found that the *Hand2* domain of expression strongly correlated inversely with that of *Gli3* (Fig. 7). This observation is compatible with Gli3R continuously repressing *Hand2* transcription, even at later stages while the early repression of *Gli3* transcription by Hand2 seems to be taken over by Hox13 products at later stages.

**Figure 7.**
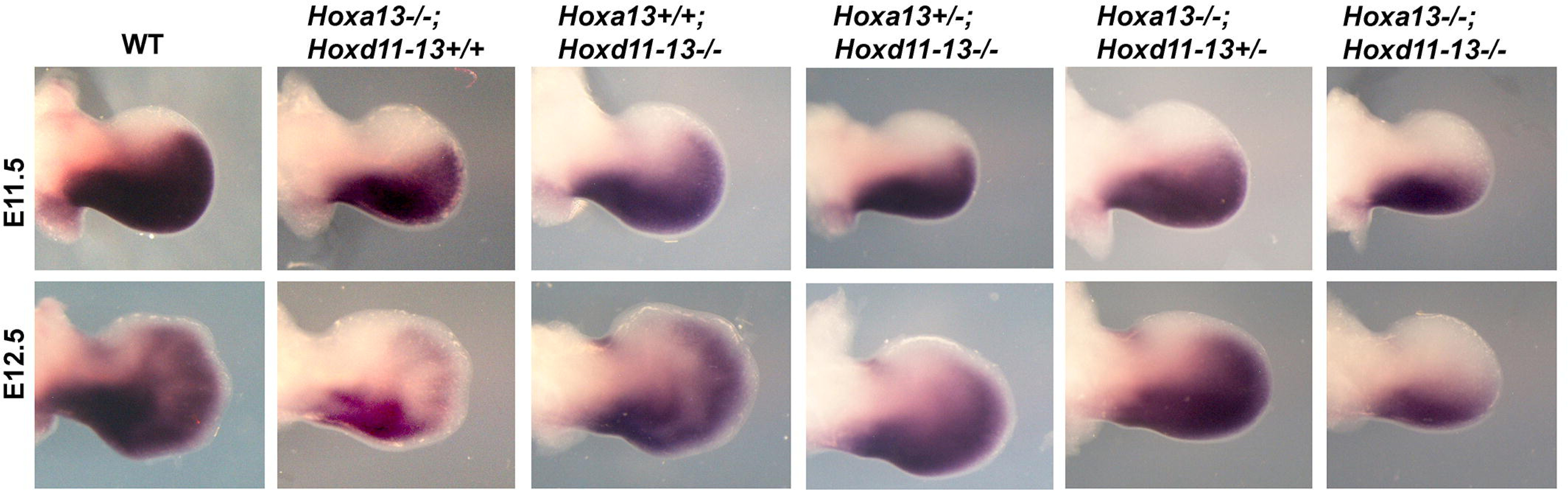
*Hand2* gene expression in *Hoxa1;Hoxd11-13* compound mutant limb buds. In situ hybridization showing the expression of *Hand2* at E11.5 and E12.5 in the genotypes of the allelic series as indicated at the figure top.

## DISCUSSION

In this study we have used a *Hoxa13* null mutant allele to further analyze the contribution of *Hox* genes to distal limb bud development. In particular, we have investigated the mechanisms underlying the loss of digit 1 in *Hoxa13* null mice. The thumb is the last digit to form (Frobisch et al., 2007) and the one with higher risk of developmental disruption. More than 1000 syndromes in the Online Mendelian Inheritance in Man (OMIM) database have hypoplastic thumbs (Oberg, 2014).

### The absence of *Hoxd13* expression in digit 1 progenitor cells explains the lack of digit 1 in *Hoxa13* mutants

Our results show that in the absence of *Hoxa13*, the domain of expression of *Hoxd13* is similar to that of *Hoxd12*, with an anterior limit coincident with the anterior border of digit 2 and therefore, with no evidence of the so-called reverse colinearity (Montavon et al., 2008; Nelson et al., 1996). This is in agreement with the crucial role of Hoxa13 in promoting the second phase of *Hoxd* gene expression and reveals that Hoxa13 is continuously required for this function (Beccari et al., 2016; Ros, 2016; Sheth et al., 2016; Sheth et al., 2014).

In the *Hoxa13* mutant, the lack of anterior propagation of *Hoxd13* expression leaves digit 1 territory devoid of Hox13 paralogues, a situation equivalent to *Hoxa13;Hoxd11-13* or *Hoxa13;Hoxd13* double mutants and that is considered sufficient to preclude the formation of the digit condensations (Fromental-Ramain et al., 1996; Sheth et al., 2012; Zakany et al., 1997). Digit patterning, the generation of a periodic digit-interdigit pattern, is under the control of a Turing-type or reaction-diffusion mechanism in which *5’Hox* genes modulate, in a dose dependent manner, the digit spacing (Sheth et al., 2012). The current model predicts that in the total absence of distal *Hox* genes (*Hoxa13, Hoxd11-13*), the area of digit patterning is strongly reduced and no distinct digital condensations form, as occurs in the digit 1 territory in the *Hoxa13* mutant.

### Loss of *Hoxa13* function is associated with a gain in Gli3R activity in the anterior mesoderm

Although Hoxa13 seems to play a role in the anterior spread of *Hoxd13*, this function is not required in the absence of *Gli3.* Actually, in the absence of *Gli3*, regardless of whether or not *Hoxa13* is present, the second phase of expression of 5’*Hoxd* genes uniformly spans the AP axis of the autopod (Litingtung et al., 2002; te Welscher et al., 2002b). Consequently, the prospective digit 1 progenitors (those located at the anterior border) form a digit with posterior identity because they express a combination of Hox products that includes Hoxd12 and Hoxd11 in addition to Hoxd13, which can be interpreted as a “transformation” or “posteriorization” of digit 1 identity.

Therefore, we reasoned that during normal development, Hoxa13 could modulate the repressor function of Gli3R to permit a fully realized second phase of *Hoxd13* expression. It has been suggested that because of its higher transcriptional efficiency, *Hoxd13* seems to be less sensitive than *Hoxd12* and *Hoxd11* to repression by Gli3R, and therefore is the only 5’Hoxd member normally expressed in digit 1, the area of maximum GLI3R level (Montavon et al., 2008). Thus, Hoxa13 could potentially attenuate Gli3R levels sufficiently to permit the spread of *Hoxd13* but not the other *5’Hoxd* genes.

Supporting this hypothesis, we provide compelling evidence of increased Gli3R activity in the anterior mesoderm of the *Hoxa13* mutants. First, the expression of bona fide Gli3R activated targets, such as *Pax9* and *EphA3* (McGlinn et al., 2005), are clearly upregulated while repressed targets, such as *Jag1* (McGlinn et al., 2005) and *Grem1* (Vokes et al., 2008) are downregulated in the anterior mutant mesoderm. The failure of *Grem1* to propagate to the anterior mesoderm also explains the mild downregulation of *Fgf8* at the anterior border, which could secondarily contribute to the *Hoxa13* phenotype. Second, we also show that both *Gli3* mRNA transcription and Gli3R protein levels are elevated in the anterior mutant mesoderm as determined by RT-PCR, RNA-seq and immunoblotting, respectively.

At first glimpse it may seem contradictory that digit 1 is lost because of excess of Gli3R as this is the only digit that reportedly forms in the hindlimb with high levels of Gli3R such as in the *Shh* and *Ozd* mutants (Chiang et al., 2001; Kraus et al., 2001; Ros et al., 2003). However, it should be noted that, as in the hindlimb, the small growth that occurs in the *Shh* null forelimb is also accompanied by expression of both *Hoxa13* and *Hoxd13*. Due to the massive cell death that occurs in both *Shh* and *Ozd* mutant limb buds, a cell lineage study would be required to determine the origin of the progenitors of the single rudimentary digit that forms.

### Hoxa13 dependent regulation of *Gli3* transcription and mutual transcriptional repression between *Gli3* and distal *Hox* genes

Interestingly, analysis of the *Gli3* expression pattern by in situ hybridization unexpectedly showed that it was highly altered in the absence of *Hoxa13* pointing to a role for Hox proteins in the transcriptional regulation of *Gli3*. Our analysis uncovered a previously unappreciated and highly dynamic pattern of *Gli3* expression that, due to similarity with that of some 5’*Hoxd* genes, we describe as evolving in two phases. After an initial wave of expression in the early limb bud mesoderm except for the most posterior *Shh*-expressing area, *Gli3* transcription becomes progressively downregulated from posterior to anterior until becoming confined to a proximal band at the zeugopod-autopod boundary. A second wave of expression is gradually established along the distal digital plate. As a consequence, two separate domains of *Gli3* expression are clearly seen in the E12.5 autopod separated by a band of tissue devoid of transcripts that corresponds to the wrist.

The dynamic pattern of *Gli3* transcription is highly altered in the absence of *Hoxa13* as the downregulation of the first wave of expression is delayed and incomplete remaining over digit 1 territory so that when the second phase of expression is established, the gap between the two phases lies between prospective digit 1 and 2. This altered pattern of expression is identical to that of *Hoxd10* and *Hoxd11* in the absence of *Hoxa13*. Indeed, the dynamics of the downregulation of the *Gli3* first phase is totally coincident with the progression of the second phase of *Hoxd13* genes, both in the wild type bud and in *Hoxa13;Hoxd11-13* allelic series, pointing to a mutual transcriptional repression between Gli3R and *Hoxd* genes and uncovering an additional level of interaction between *Hox* genes and the Shh/Gli3 pathway (Fig. 8). This is supported by the presence of several Hox13 binding sites in the *Gli3* genomic landscape the function of which definitely deserves further investigation.

**Figure 8.**
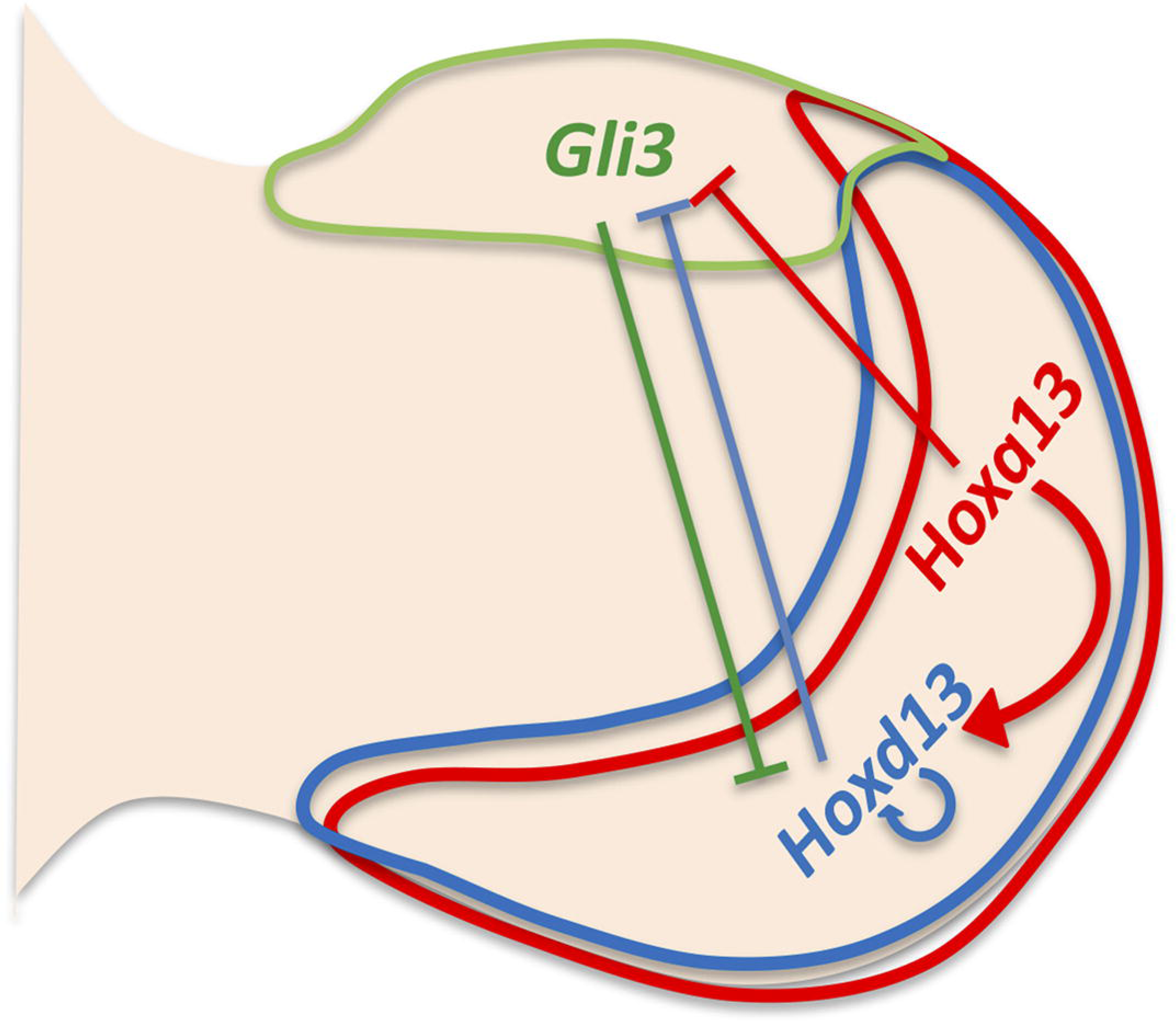
Schematic diagram indicating the interactions between *Gli3* and *5’Hox* genes.

### No evidence of Hoxa13-Gli3 physical interaction

We have also considered the possibility that Hoxa13 interacts with Gli3R at the protein level, as has been shown for Hoxd12 and Hoxd13 (Chen et al., 2004), and that this binding modifies GLI3R activity either by sequestering it or transforming its repressor activity into an activator. However, coimmunoprecipitation (CoIP) of E 11.5 limb bud lysates using antibodies specific to HOX13 and GLI3 (Chen et al., 2004; Knosp et al., 2004; Wen et al., 2010) didn’t detect such interaction (authors unpublished results). It is noteworthy that a recent screen based on affinity purification of endogenous protein and mass spectrometry (RIME, Mohammed et al., 2016) did not identify GLI3 as an interacting partner of HOXA13 (Marie Kmita, personal communication) supporting the conclusion that the gain in Gli3R activity observed in the anterior autopod of *Hoxa13* mutants is primarily the result of the transcriptional control exerted by Hox13 proteins.

### Summary

It is well known that Gli3R regulates the transcription of the *5’Hox* genes in the anterior mesoderm in a dose-dependent manner (Litingtung et al., 2002; te Welscher et al., 2002b); Supplementary Fig. 2). Gli3R repressor activity may be mediated by direct binding of Gli3R to the enhancers in the regulatory *Hoxd* landscape (Vokes et al., 2008). Here we provide compelling evidence for the implication of Hoxa13 in the attenuation of Gli3R expression required for *Hoxd13* transcriptional spread into the most anterior mesoderm (reverse collinearity). Hoxa13 together with Hoxd13 repress *Gli3* transcription establishing a mutual antagonism necessary for the correct anterior-posterior asymmetry of the autopod. The acquisition of the repressive activity of distal Hox products on *Gli3* transcription may have been determinant in the reduction of the anterior skeletal elements and digits that gradually occurred during the fin-to-limb transition (Sheth et al., 2012; Onimaru et al., 2015)

## MATERIAL AND METHODS

### Embryos

The *Hoxa13* (Fromental-Ramain et al., 1996), *Hoxd*^*Del(11–13)*^ (Zakany and Duboule, 1996) and *Gli3* (*Gli3*^*XtJ*^ Jackson allele; (Hui and Joyner, 1993)) mutant lines were kindly provided by Pierre Chambon, Denis Duboule and Rolf Zeller, respectively. The *Hoxa13* line was maintained on a C57BL/6 genetic background and the *Gli3* line on a mixed CD1 and C57BL/6 genetic background. Genotyping was performed using tail biopsies or embryonic membranes according to previously published reports. Embryos of the desired embryonic embryonic day (E) were obtained by cesarean section. All animal procedures were conducted accordingly to the EU regulations and 3R principles and reviewed and approved by the Bioethics Committee of the University of Cantabria, and according to the ethical guidelines of the Institutional Animal Care and Use Committee (IACUC) at NCI-Frederick under protocol #ASP-12-405.

### Skeletal Preparations and *in situ* hybridization

Whole-mount skeletal preparations were performed by staining with Alcian blue 8GX (Sigma Aldrich) and Alizarin red S (Sigma Aldrich) following standard protocols. Briefly, the specimens were fixed in 95% ethanol, skinned and eviscerated before staining, cleared in a series of KOH solutions and stored in glycerol. Whole mount *in situ* hybridization was performed according to standard procedures using digoxigenin labeled antisense riboprobes. At least 2 embryos per stage and genotype were analyzed. The probes used were *Sox9, Hoxd13, Hoxd12, Hoxd11, Hoxd10, Hoxd9, Hoxd4, Hoxa11, Gli3, Pax9, Jagg1, EphA3, Hand*1, *Hand2, Bmp4, Shh, Gli1, Ptc1, Dusp6, Spry4, Grem1* and *Fgf8.*

### Western Blot

For immunoblot (Western blot) analysis, dissected autopod regions of E11.5 embryos were lysed with ice-cold RIPA buffer. 10% sodium dodecyl sulfate-polyacrylamide gel electrophoresis was used to resolve Gli3-190 from Gli3-83 protein, and the mouse monoclonal Gli3 clone 6F5 anti-Gli3-N antibody was used (kindly provided by Dr. Scales at Genentech; Wen et al., 2010). βactin (mouse monoclonal C4SC-4778, Santa Cruz) was assessed as control for normalization. Three independent experiments were performed.

### Quantitative real-time PCR (qRT-PCR)

The region of digit 1 (Fig. 4E) was dissected in cold RNAse-free PBS from E11.75 wild type and *Hoxa13* null embryos. Total RNA was extracted with RNeasy® Plus Micro Kit (Qiagen) and 50 ng of total RNA was reverse transcribed to produce first-strand cDNA with iScript™ cDNA Synthesis kit (Bio-Rad) using standard conditions. qRT-PCR was carried out on an Applied Biosystems StepOnePlus™ using SYBR Green Supermix (Bio-Rad) and the data were analyzed using the StepOne(tm) software. The primers used to amplify *Gli3* and *Hoxd13* were previously described in (Huang et al., 2016). Relative transcript levels were normalized to *Vimentin* (Huang et al., 2016). Four biological replicates were analyzed for each genotype, with at least two technical replicates for each sample. The expression levels of mutant samples were calculated relative to wild-type controls (average set to 100%). The significance of all differences was assessed using Student-test, being statistically significant when p-0.05. GraphPad Prism5.0 (LaJolla, CA) was used for graph and statistics analysis. Histogram bars represent the average expression values after normalization to Vimentin (standard deviation shown as error bars).

## Acknowledgments

We thank Genentech for the 6F5GLI3 antibody and Laura Galán, Mar Rodriguez and Víctor Campa for excellent technical assistance. We are most grateful to Berta Casar, Lorena Agudo and Endika Haro for helpful discussions.

## Author contributions

MF Bastida: Conceptualization; Investigation; Validation; Writing—review and editing R Perez-Gomez: Formal analysis; Investigation; Methodology

A Trofka: Formal analysis; Investigation; Methodology

R Sheth: Conceptualization; Formal analysis; Investigation; Writing—review and editing

HS Stadler: Formal analysis; Investigation; Funding acquisition; Writing—review and editing

S Mackem: Conceptualization; Formal analysis; Investigation; Funding acquisition; Writing—review and editing

MA Ros: Conceptualization; Formal analysis; Investigation; Funding acquisition; Supervision; Project administration; Writing—original draft

---

**Supplementary Figure 1**

**Altered *Hoxd* gene expression in *Hoxa13* mutant limb buds.**

Forelimb autopods of wild type and *Hoxa13* homozygous mutants hybridized with *Hoxd4, Hoxd9, Hoxd10* and *Hoxa11* at E 12.5. The arrows point to altered expression patterns in homozygous mutants.

**Supplementary Figure 2**

**Expression of markers of the anterior mesoderm in *Hoxa13* mutant limb buds.**

Pattern of expression of *Tbx2* (**A**), *Tbx3* (**B**) and *Alx4* (**C**) in E11.5 forelimb buds of wild type and *Hoxa13* homozygous mutants. No obvious modification in the expression patterns is observed in the mutant.

**Supplementary Figure 3**

***Hox* gene expression in limb buds of the *Hoxa13;Gli3* allelic series.**

E12.5 forelimb buds are shown with the genotype indicated at the top and the hybridization probe on the left. Note that in the absence of *Gli3*, disregarding the presence or not of *Hoxa13, Hoxd13* expression spans the entire autopod.

**Supplementary Figure 4**

**Hoxa13 protein levels increase with reduction in Hoxd11-13 dosage.**

1% SDS lysates of dissected distal limb bud (E12.5 autopod region) from wildtype and *Hoxd11-13* mutants were electrophoresed, blotted, and probed with affinity-purified polyclonal anti-Hoxd13 antibody which recognizes both Hoxa13 and Hoxd13 (Chen et al., 2004), and with anti-Vinculin (1:1,000, Sigma # V9264). Bands were visualized with fluorescent secondary antibodies and quantified using the Odyssey Li-Cor system. Hox13 fluorescence signals were normalized to Vinculin, and three independent samples were analyzed for each genotype. Significance of differences were determined using the two-tailed, Student’s t-test. An approximately 7-fold increase in Hoxa13 expression in *Hoxd11-13*-/- limb buds compared with wildtype was significant at p=0.01 and a 2-fold increase in *Hoxd11-13*+/- was significant at p=0.05.

## REFERENCES

Ahn, S. and Joyner, A. L. (2004). Dynamic changes in the response of cells to positive hedgehog signaling during mouse limb patterning. Cell 118, 505–16.

Andrey, G., Montavon, T., Mascrez, B., Gonzalez, F., Noordermeer, D., Leleu, M., Trono, D., Spitz, F. and Duboule, D. (2013). A switch between topological domains underlies HoxD genes collinearity in mouse limbs. Science 340, 1234167.

Bastida, M. F., Delgado, M. D., Wang, B., Fallon, J. F., Fernandez-Teran, M. and Ros, M. A. (2004). Levels of Gli3 repressor correlate with Bmp4 expression and apoptosis during limb development. Dev Dyn 231, 148–60.

Bastida, M. F., Sheth, R. and Ros, M. A. (2009). A BMP-Shh negative-feedback loop restricts Shh expression during limb development. Development 136, 3779–89.

Beccari, L., Yakushiji-Kaminatsui, N., Woltering, J. M., Necsulea, A., Lonfat, N., Rodriguez-Carballo, E., Mascrez, B., Yamamoto, S., Kuroiwa, A. and Duboule, D. (2016). A role for HOX13 proteins in the regulatory switch between TADs at the HoxD locus. Genes Dev 30, 1172–86.

Benazet, J. D., Bischofberger, M., Tiecke, E., Goncalves, A., Martin, J. F., Zuniga, A., Naef, F. and Zeller, R. (2009). A self-regulatory system of interlinked signaling feedback loops controls mouse limb patterning. Science 323, 1050–3.

Berlivet, S., Paquette, D., Dumouchel, A., Langlais, D., Dostie, J. and Kmita, M. (2013). Clustering of tissue-specific sub-TADs accompanies the regulation of HoxA genes in developing limbs. PLoS Genet 9, e1004018.

Boulet, A. M. and Capecchi, M. R. (2004). Multiple roles of Hoxa11 and Hoxd11 in the formation of the mammalian forelimb zeugopod. Development 131, 299–309.

Burke, A. C., Nelson, C. E., Morgan, B. A. and Tabin, C. (1995). Hox genes and the evolution of vertebrate axial morphology. Development 121, 333–46.

Capellini, T. D., Di Giacomo, G., Salsi, V., Brendolan, A., Ferretti, E., Srivastava, D., Zappavigna, V. and Selleri, L. (2006). Pbx1/Pbx2 requirement for distal limb patterning is mediated by the hierarchical control of Hox gene spatial distribution and Shh expression. Development 133, 2263–73.

Chen, Y., Knezevic, V., Ervin, V., Hutson, R., Ward, Y. and Mackem, S. (2004). Direct interaction with Hoxd proteins reverses Gli3-repressor function to promote digit formation downstream of Shh. Development 131, 2339–47.

Chiang, C., Litingtung, Y., Harris, M. P., Simandl, B. K., Li, Y., Beachy, P. A. and Fallon, J. F. (2001). Manifestation of the limb prepattern: limb development in the absence of sonic hedgehog function. Dev Biol 236, 421–35.

Davis, A. P., Witte, D. P., Hsieh-Li, H. M., Potter, S. S. and Capecchi, M. R. (1995). Absence of radius and ulna in mice lacking hoxa-11 and hoxd-11. Nature 375, 791–5.

Dixon, J. R., Selvaraj, S., Yue, F., Kim, A., Li, Y., Shen, Y., Hu, M., Liu, J. S. and Ren, B. (2012). Topological domains in mammalian genomes identified by analysis of chromatin interactions. Nature 485, 376–80.

Dolle, P., Dierich, A., LeMeur, M., Schimmang, T., Schuhbaur, B., Chambon, P. and Duboule, D. (1993). Disruption of the Hoxd-13 gene induces localized heterochrony leading to mice with neotenic limbs. Cell 75, 431–41.

Fernandez-Teran, M., Piedra, M. E., Rodriguez-Rey, J. C., Talamillo, A. and Ros, M. A. (2003). Expression and regulation of eHAND during limb development. Dev Dyn 226, 690–701.

Frobisch, N. B., Carroll, R. L. and Schoch, R. R. (2007). Limb ossification in the Paleozoic branchiosaurid Apateon (Temnospondyli) and the early evolution of preaxial dominance in tetrapod limb development. Evol Dev 9, 69–75.

Fromental-Ramain, C., Warot, X., Messadecq, N., LeMeur, M., Dolle, P. and Chambon, P. (1996). Hoxa-13 and Hoxd-13 play a crucial role in the patterning of the limb autopod. Development 122, 2997–3011.

Galli, A., Robay, D., Osterwalder, M., Bao, X., Benazet, J. D., Tariq, M., Paro, R., Mackem, S. and Zeller, R. (2010). Distinct roles of Hand2 in initiating polarity and posterior Shh expression during the onset of mouse limb bud development. PLoS Genet 6, e1000901.

Goodman, F. R., Bacchelli, C., Brady, A. F., Brueton, L. A., Fryns, J. P., Mortlock, D. P., Innis, J. W., Holmes, L. B., Donnenfeld, A. E., Feingold, M. et al. (2000). Novel HOXA13 mutations and the phenotypic spectrum of hand-foot-genital syndrome. Am J Hum Genet 67, 197–202.

Harfe, B. D., Scherz, P. J., Nissim, S., Tian, H., McMahon, A. P. and Tabin, C. J. (2004). Evidence for an expansion-based temporal Shh gradient in specifying vertebrate digit identities. Cell 118, 517–28.

Huang, B. L., Trofka, A., Furusawa, A., Norrie, J. L., Rabinowitz, A. H., Vokes, S. A., Mark Taketo, M., Zakany, J. and Mackem, S. (2016). An interdigit signalling centre instructs coordinate phalanx-joint formation governed by 5’Hoxd-Gli3 antagonism. Nat Commun 7, 12903.

Hui, C. C. and Joyner, A. L. (1993). A mouse model of greig cephalopolysyndactyly syndrome: the extra-toesJ mutation contains an intragenic deletion of the Gli3 gene. Nat Genet 3, 241–6.

Innis, J. W., Goodman, F. R., Bacchelli, C., Williams, T. M., Mortlock, D. P., Sateesh, P., Scambler, P. J., McKinnon, W. and Guttmacher, A. E. (2002). A HOXA13 allele with a missense mutation in the homeobox and a dinucleotide deletion in the promoter underlies Guttmacher syndrome. Hum Mutat 19, 573–4.

Kherdjemil, Y., Lalonde, R. L., Sheth, R., Dumouchel, A., de Martino, G., Pineault, K. M., Wellik, D. M., Stadler, H. S., Akimenko, M. A. and Kmita, M. (2016). Evolution of Hoxa11 regulation in vertebrates is linked to the pentadactyl state. Nature 539, 89–92.

Khokha, M. K., Hsu, D., Brunet, L. J., Dionne, M. S. and Harland, R. M. (2003). Gremlin is the BMP antagonist required for maintenance of Shh and Fgf signals during limb patterning. Nat Genet 34, 303–7.

Kmita, M., Fraudeau, N., Herault, Y. and Duboule, D. (2002). Serial deletions and duplications suggest a mechanism for the collinearity of Hoxd genes in limbs. Nature 420, 145–50.

Kmita, M., Tarchini, B., Zakany, J., Logan, M., Tabin, C. J. and Duboule, D. (2005). Early developmental arrest of mammalian limbs lacking HoxA/HoxD gene function. Nature 435, 1113–6.

Knosp, W. M., Saneyoshi, C., Shou, S., Bachinger, H. P. and Stadler, H. S. (2007). Elucidation, quantitative refinement, and in vivo utilization of the HOXA13 DNA binding site. J Biol Chem 282, 6843–53.

Knosp, W. M., Scott, V., Bachinger, H. P. and Stadler, H. S. (2004). HOXA13 regulates the expression of bone morphogenetic proteins 2 and 7 to control distal limb morphogenesis. Development 131, 4581–92.

Kraus, P., Fraidenraich, D. and Loomis, C. A. (2001). Some distal limb structures develop in mice lacking Sonic hedgehog signaling. Mech Dev 100, 45–58.

Laufer, E., Nelson, C. E., Johnson, R. L., Morgan, B. A. and Tabin, C. (1994). Sonic hedgehog and Fgf-4 act through a signaling cascade and feedback loop to integrate growth and patterning of the developing limb bud. Cell 79, 993–1003.

Lewandowski, J. P., Du, F., Zhang, S., Powell, M. B., Falkenstein, K. N., Ji, H. and Vokes, S. A. (2015). Spatiotemporal regulation of GLI target genes in the mammalian limb bud. Dev Biol 406, 92–103.

Litingtung, Y., Dahn, R. D., Li, Y., Fallon, J. F. and Chiang, C. (2002). Shh and Gli3 are dispensable for limb skeleton formation but regulate digit number and identity. Nature 418, 979–83.

Lonfat, N., Montavon, T., Darbellay, F., Gitto, S. and Duboule, D. (2014). Convergent evolution of complex regulatory landscapes and pleiotropy at Hox loci. Science 346, 1004–6.

McGlinn, E., van Bueren, K. L., Fiorenza, S., Mo, R., Poh, A. M., Forrest, A., Soares, M. B., Bonaldo Mde, F., Grimmond, S., Hui, C. C. et al. (2005). Pax9 and Jagged1 act downstream of Gli3 in vertebrate limb development. Mech Dev 122, 1218–33.

Michos, O., Panman, L., Vintersten, K., Beier, K., Zeller, R. and Zuniga, A. (2004). Gremlin-mediated BMP antagonism induces the epithelial-mesenchymal feedback signaling controlling metanephric kidney and limb organogenesis. Development 131, 3401–10.

Minowada, G., Jarvis, L. A., Chi, C. L., Neubuser, A., Sun, X., Hacohen, N., Krasnow, M. A. and Martin, G. R. (1999). Vertebrate Sprouty genes are induced by FGF signaling and can cause chondrodysplasia when overexpressed. Development 126, 4465–75.

Mitsubuchi, H. and Endo, F. (2006). [Hand-foot-genital syndrome]. Nihon Rinsho Suppl 2, 647–8.

Montavon, T. and Duboule, D. (2013). Chromatin organization and global regulation of Hox gene clusters. Philos Trans R Soc Lond B Biol Sci 368, 20120367.

Montavon, T., Le Garrec, J. F., Kerszberg, M. and Duboule, D. (2008). Modeling Hox gene regulation in digits: reverse collinearity and the molecular origin of thumbness. Genes Dev 22, 346–59.

Montavon, T., Soshnikova, N., Mascrez, B., Joye, E., Thevenet, L., Splinter, E., de Laat, W., Spitz, F. and Duboule, D. (2011). A regulatory archipelago controls Hox genes transcription in digits. Cell 147, 1132–45.

Nelson, C. E., Morgan, B. A., Burke, A. C., Laufer, E., DiMambro, E., Murtaugh, L. C., Gonzales, E., Tessarollo, L., Parada, L. F. and Tabin, C. (1996). Analysis of Hox gene expression in the chick limb bud. Development 122, 1449–66.

Niswander, L., Jeffrey, S., Martin, G. R. and Tickle, C. (1994). A positive feedback loop coordinates growth and patterning in the vertebrate limb. Nature 371, 609–12.

Oberg, K. C. (2014). Review of the molecular development of the thumb: digit primera. Clin Orthop Relat Res 472, 1101–5.

Onimaru, K., Kuraku, S., Takagi, W., Hyodo, S., Sharpe, J. and Tanaka, M. (2015). A shift in anterior-posterior positional information underlies the fin-to-limb evolution. Elife 4.

Osterwalder, M., Barozzi, I., Tissieres, V., Fukuda-Yuzawa, Y., Mannion, B. J., Afzal, S. Y., Lee, E. A., Zhu, Y., Plajzer-Frick, I., Pickle, C. S. et al. (2018). Enhancer redundancy provides phenotypic robustness in mammalian development. Nature 554, 239–243.

Osterwalder, M., Speziale, D., Shoukry, M., Mohan, R., Ivanek, R., Kohler, M., Beisel, C., Wen, X., Scales, S. J., Christoffels, V. M. et al. (2014). HAND2 targets define a network of transcriptional regulators that compartmentalize the early limb bud mesenchyme. Dev Cell 31, 345–57.

Perez, W. D., Weller, C. R., Shou, S. and Stadler, H. S. (2010). Survival of Hoxa13 homozygous mutants reveals a novel role in digit patterning and appendicular skeletal development. Dev Dyn 239, 446–57.

Ros, M. A. (2016). HOX13 proteins: the molecular switcher in Hoxd bimodal regulation. Genes Dev 30, 1135–7.

Ros, M. A., Dahn, R. D., Fernandez-Teran, M., Rashka, K., Caruccio, N. C., Hasso, S. M., Bitgood, J. J., Lancman, J. J. and Fallon, J. F. (2003). The chick oligozeugodactyly (ozd) mutant lacks sonic hedgehog function in the limb. Development 130, 527–37.

Scotti, M. and Kmita, M. (2012). Recruitment of 5’ Hoxa genes in the allantois is essential for proper extra-embryonic function in placental mammals. Development 139, 731–9.

Scherz, P. J., Harfe, B. D., McMahon, A. P. and Tabin, C. J. (2004). The limb bud Shh-Fgf feedback loop is terminated by expansion of former ZPA cells. Science 305, 396–9.

Shaut, C. A., Keene, D. R., Sorensen, L. K., Li, D. Y. and Stadler, H. S. (2008). HOXA13 Is essential for placental vascular patterning and labyrinth endothelial specification. PLoS Genet 4, e1000073.

Sheth, R., Barozzi, I., Langlais, D., Osterwalder, M., Nemec, S., Carlson, H. L., Stadler, H. S., Visel, A., Drouin, J. and Kmita, M. (2016). Distal Limb Patterning Requires Modulation of cis-Regulatory Activities by HOX13. Cell Rep 17, 2913–2926.

Sheth, R., Bastida, M. F., Kmita, M. and Ros, M. (2014). “Self-regulation,” a new facet of Hox genes’ function. Dev Dyn 243, 182–91.

Sheth, R., Gregoire, D., Dumouchel, A., Scotti, M., Pham, J. M., Nemec, S., Bastida, M. F., Ros, M. A. and Kmita, M. (2013). Decoupling the function of Hox and Shh in developing limb reveals multiple inputs of Hox genes on limb growth. Development 140, 2130–8.

Sheth, R., Marcon, L., Bastida, M. F., Junco, M., Quintana, L., Dahn, R., Kmita, M., Sharpe, J. and Ros, M. A. (2012). Hox genes regulate digit patterning by controlling the wavelength of a Turing-type mechanism. Science 338, 1476–80.

Smith, T. G., Karlsson, M., Lunn, J. S., Eblaghie, M. C., Keenan, I. D., Farrell, E. R., Tickle, C., Storey, K. G. and Keyse, S. M. (2006). Negative feedback predominates over cross-regulation to control ERK MAPK activity in response to FGF signalling in embryos. FEBS Lett. 2006 Jul 24 580(17), 4242–5.

Spitz, F., Gonzalez, F., Peichel, C., Vogt, T. F., Duboule, D. and Zakany, J. (2001). Large scale transgenic and cluster deletion analysis of the HoxD complex separate an ancestral regulatory module from evolutionary innovations. Genes Dev 15, 2209–14.

Stadler, H. S., Higgins, K. M. and Capecchi, M. R. (2001). Loss of Eph-receptor expression correlates with loss of cell adhesion and chondrogenic capacity in Hoxa13 mutant limbs. Development 128, 4177–88.

Tabin, C. and Wolpert, L. (2007). Rethinking the proximodistal axis of the vertebrate limb in the molecular era. Genes Dev 21, 1433–42.

Tarchini, B. and Duboule, D. (2006). Control of Hoxd genes’ collinearity during early limb development. Dev Cell 10, 93–103.

te Welscher, P., Fernandez-Teran, M., Ros, M. A. and Zeller, R. (2002a). Mutual genetic antagonism involving GLI3 and dHAND prepatterns the vertebrate limb bud mesenchyme prior to SHH signaling. Genes Dev 16, 421–6.

te Welscher, P., Zuniga, A., Kuijper, S., Drenth, T., Goedemans, H. J., Meijlink, F. and Zeller, R. (2002b). Progression of vertebrate limb development through SHH-mediated counteraction of GLI3. Science 298, 827–30.

Verheyden, J. M. and Sun, X. (2008). An Fgf/Gremlin inhibitory feedback loop triggers termination of limb bud outgrowth. Nature 454, 638–41.

Vokes, S. A., Ji, H., Wong, W. H. and McMahon, A. P. (2008). A genome-scale analysis of the cis-regulatory circuitry underlying sonic hedgehog-mediated patterning of the mammalian limb. Genes Dev 22, 2651–63.

Wang, B., Fallon, J. F. and Beachy, P. A. (2000). Hedgehog-regulated processing of Gli3 produces an anterior/posterior repressor gradient in the developing vertebrate limb. Cell 100, 423–34.

Wen, X., Lai, C. K., Evangelista, M., Hongo, J. A., de Sauvage, F. J. and Scales, S. J. (2010). Kinetics of hedgehog-dependent full-length Gli3 accumulation in primary cilia and subsequent degradation. Mol Cell Biol 30, 1910–22.

Williams, T. M., Williams, M. E., Heaton, J. H., Gelehrter, T. D. and Innis, J. W. (2005). Group 13 HOX proteins interact with the MH2 domain of R-Smads and modulate Smad transcriptional activation functions independent of HOX DNA-binding capability. Nucleic Acids Res 33, 4475–84.

Woltering, J. M. and Duboule, D. (2010). The origin of digits: expression patterns versus regulatory mechanisms. Dev Cell 18, 526–32.

Woltering, J. M., Noordermeer, D., Leleu, M. and Duboule, D. (2014). Conservation and divergence of regulatory strategies at Hox Loci and the origin of tetrapod digits. PLoS Biol 12, e1001773.

Zakany, J. and Duboule, D. (1996). Synpolydactyly in mice with a targeted deficiency in the HoxD complex. Nature 384, 69–71.

Zakany, J. and Duboule, D. (2007). The role of Hox genes during vertebrate limb development. Curr Opin Genet Dev 17, 359–66.

Zakany, J., Fromental-Ramain, C., Warot, X. and Duboule, D. (1997). Regulation of number and size of digits by posterior Hox genes: a dose-dependent mechanism with potential evolutionary implications. Proc Natl Acad Sci U S A 94, 13695–700.

Zeller, R., Lopez-Rios, J. and Zuniga, A. (2009). Vertebrate limb bud development: moving towards integrative analysis of organogenesis. Nat Rev Genet 10, 845–58.

Zuniga, A. (2015). Next generation limb development and evolution: old questions, new perspectives. Development 142, 3810–20.

